# Development of the preterm infant gut and gastric residuals microbiome

**DOI:** 10.1101/2025.06.21.660859

**Authors:** Nadav Moriel, Leah Jones, Esty Harpanes, Nechama Rakow, Shimrit Shmorak, Smadar Eventov Friedman, Noa Ofek Shlomai, Moran Yassour

## Abstract

Prematurity, defined as birth before 37 completed weeks of gestation, is the leading cause of mortality in children under five and affects approximately 11% of live births globally—around 15 million infants each year. Despite advances in neonatal care, preterm infants remain at an increased risk for a range of complications. One widely used clinical practice in neonatal intensive care units (NICUs) is the monitoring of gastric residuals (GRs) to assess feeding tolerance and guide enteral nutrition. While the clinical significance of GRs is debated, their microbial composition has not been extensively studied.

In this study, we performed metagenomic sequencing of 199 stool and 69 GR samples from 39 preterm infants during hospitalization to characterize their gut and stomach microbiomes. To our knowledge, this is the first study to describe the microbial landscape of GR in preterm infants. We identified 11 distinct GR clusters, often dominated by *Staphylococcus*, *Streptococcus*, and *Klebsiella*, with microbial diversity correlating with GR aspiration frequency. Longitudinal analysis revealed temporal colonization patterns, with early dominance of *Staphylococcus epidermidis* and *Bradyrhizobium*, and later emergence of *Escherichia coli*, *Staphylococcus hominis*, and *Streptococcus thermophilus*.

In stool samples, 8 microbial clusters were found, frequently enriched with *Enterobacteriaceae*. Early samples rich in *S. epidermidis* were associated with higher gestational age and lower microbial richness, while *Bifidobacterium breve*, a beneficial gut commensal, appeared later in hospitalization. Comparative analysis showed overlap between gut and gastric microbiota, though stomach samples were more dynamic and exhibited less intra-subject similarity.

Strain-level resolution revealed both subject-specific (e.g., *E. coli*, *K. pneumoniae*) and widely shared (e.g., *S. epidermidis*) taxa. We also identified a pathogenic *Klebsiella aerogenes* strain associated with bacteremia with distinct genomic features a week ahead of its first clinical isolation. These findings provide novel insights into the dynamic and niche-specific microbial colonization of preterm infants.

## Introduction

The gastrointestinal tract is heavily colonized by trillions of microorganisms known as the gut microbiota, which critically support host physiology, healthy development, and function^1–4^. Inoculation of the gut microbiome of newborns begins at birth, and the neonatal gut environment and maturity impact its composition^5^. Establishment of the gut barrier leads to a shift in microbial composition from the facultative anaerobes (such as *Enterobactericea* and *Enterococci*) to strict anaerobes (such as *Bacteroides*, *Bifidobacterium,* and *Clostridium*), reflecting an adaptation to the decreasing oxygen levels within the gut lumen^6,7^. Changes in the composition of the gut microbiota (dysbiosis) have been associated with a wide array of diseases, including inflammatory, autoimmune, neurological, and metabolic disorders^8–13^. Intriguingly, gut microbial dysbiosis during neonatal development is linked to disease development later in life, such as allergy, asthma, and autoimmune disorders^14–16^. Furthermore, bacteria residing in an infant’s gut are pivotal in allowing the infant to maximize absorption from their diet^17^. One of the bacteria that has been shown to benefit infants and be associated with a healthy infant gut microbial environment is *Bifidobacterium*^18–24^. Infants hospitalized in the neonatal intensive care unit (NICU) are not provided with various milk formulations^25^ and probiotics^19,21^, containing *Bifidobacterium*, despite its considerable impact on shaping the gut microbiome, due to health ministry policy^19,26,27^.

Preterm birth affects approximately 15 million infants annually, representing approximately 11% of live births worldwide, with incidence rates continuing to rise^28^. Although advances in neonatology have enhanced survival rates among the most premature and critically ill infants, these populations remain at increased risk for a broad spectrum of long-term adverse health outcomes^28^. Neonatal morbidities of preterm infants include respiratory morbidities such as respiratory distress syndrome, later leading to bronchopulmonary dysplasia, neurological morbidity including intraventricular hemorrhage and periventricular leukomalacia, retinopathy of prematurity, early and late onset sepsis, necrotizing enterocolitis, and more^29^.

Necrotizing enterocolitis (NEC) is the leading gastrointestinal cause of morbidity and mortality in neonatal intensive care units (NICUs), primarily affecting very-low-birth-weight (VLBW) premature infants (birth weight < 1500 g), with a prevalence of 5% to 10% and a mortality rate of 25% to 30%^30^. NEC is a result of a multifactorial process that includes reduced blood flow to an immature gut, (including disruptions to early immunological maturation)^31,32^. This may lead to increased gut permeability^33,34^. Changes to the gut wall that follow bacterial dysbiosis allow for the translocation of bacteria from the gut lumen into the bloodstream, potentially causing bacteremia and sepsis^35–37^. Prevention of life-threatening conditions in preterm infants often incorporates multiple clinical interventions, including prolonged intensive care monitoring and antibiotic administration, which can alter the developing microbiome.

Repetitive examination of gastric residuals (GR), collected by aspirating the gastric content of preterm infants, is a common practice performed in the NICU of Hadassah Medical Center that helps assess infant health status and guides the initiation and progression of enteral feeding in these infants^38,39^. Gastric residue monitoring aims to use abnormalities or increases in gastric residual volumes as early predictors of feeding intolerance, potentially reflecting NEC^40–42^. However, recent studies suggest that in the absence of clinical signs of feeding intolerance, avoiding pre-feeding monitoring of gastric residues is safe, and does not increase the risk for NEC, furthermore there have been reports that routine monitoring of GR does not improve clinical outcomes and may delay advancement of enteral nutrition and weight gain^43–46^.

Anatomically, the stomach and gut are interconnected segments within the seamless continuum of the gastrointestinal tract, presenting different settings for microbial colonization owing to the contrasting environmental conditions, such as gastric acidity and motility. Bacteria may overcome these environmental dissimilarities and utilize the common tract connecting the two organs to translocate from one niche to another. A study using 16S sequencing to profile the gastric microbiome reported the presence of *Bacteroides spp.* and *Bifidobacterium spp.* within this niche^47^. To date, the gastric microbiome of premature infants remains largely unexplored, particularly in terms of comprehensive characterization using metagenomic sequencing approaches.

To expand the knowledge of the GR and stool microbiome development in premature infants, we performed metagenomic sequencing of 69 GR samples and 199 samples from 39 subjects and profiled their microbial composition throughout their postnatal hospitalization. We identified composition-derived clusters within the samples and carried out comparisons involving both microbial and clinical metrics. Our results support and highlight some of the gut microbiome developmental changes occurring throughout its maturation process, such as the transition from *Staphylococcus epidermidis-*dominated communities and the chronological emergence of *Bifidobacterium-*dominated bacterial populations. Among the gastric microbiome, we found co-occurrence of bacteria also present across stool samples, including members of the *Klebsiella* and *Staphylococcus* genera. Lastly, using sequenced data, we identified a strain of *Klebsiella aerogenes,* ultimately associated with bacteremia a week before its initial detection within a blood sample, highlighting the potential of sequencing methods to provide early detection of the presence of pathogenic strains, preceding future systemic infections.

## Results

### Study Design

Premature infants, born before completing 37 weeks of gestation, are initially colonized with microbes within the first days of life^48–52^. These acquired microbial populations, consisting of bacteria, fungi, and viruses, play a pivotal role in the infant’s development and have been associated with various health conditions^53–55^. During their postnatal hospitalization, infants frequently undergo a wide spectrum of medical interventions and tests that have the potential to alter the microbial profile, such as antibiotic treatment^56–58^. Observing the establishment and dynamics of the microbiome in preterm infants, in correlation with their respective clinical status, has the potential to help understand the relationship between the host microbiome and health. An interesting clinical practice carried out in some neonatal intensive care units (NICUs) is the monitoring of gastric residual (GR) samples, which consists of aspiration of stomach content before feeding, as well as in cases of clinical indications^40^. To the best of our knowledge, this is the first study to characterize the microbial communities of the gastric residuals in premature infants using metagenomic sequencing methods.

To pursue these challenges, we collaborated with the NICU of Hadassah Ein Kerem Medical Center, and collected samples from premature infants during their post-delivery hospitalization (**Figure 1**). A total 199 stool and 69 GR samples were collected from 39 premature infants (median GA 29 weeks, mean GA 29 weeks + 1 day) and sequenced using metagenomic sequencing (6,273,587 mean read coverage). Subjects were randomized into a high or low frequency gastric residuals collection group consisting of 19 and 20 participants, respectively **(Methods)**. While all subjects underwent gastric residuals aspiration upon clinical indication, gastric residuals were routinely aspirated prior to feeding in infants within the high-frequency group. Necrotizing enterocolitis occurred in five infants (3 and 2 subjects from the low and high-frequency groups, respectively). Five premature infants, some of whom developed NEC, died during their hospitalization period (also, 3 and 2 cases from the low and high-frequency study groups, respectively). Averages of clinical metrics across the study groups were similar, including length of hospitalization (60.2 days vs. 50.2 days; p value = 0.28), birth weight z-score (0.049 vs. -0.548; pvalue = 0.059), delta z-score of discharge weight (1.44 vs. 1.15; p value = 0.24), gestational age (28w+5d vs. 29w+4d; pvalue = 0.35) and birthweight (1170 grams vs. 1130 grams; p value = 0.76; **Table 1**).

**Figure 1.**
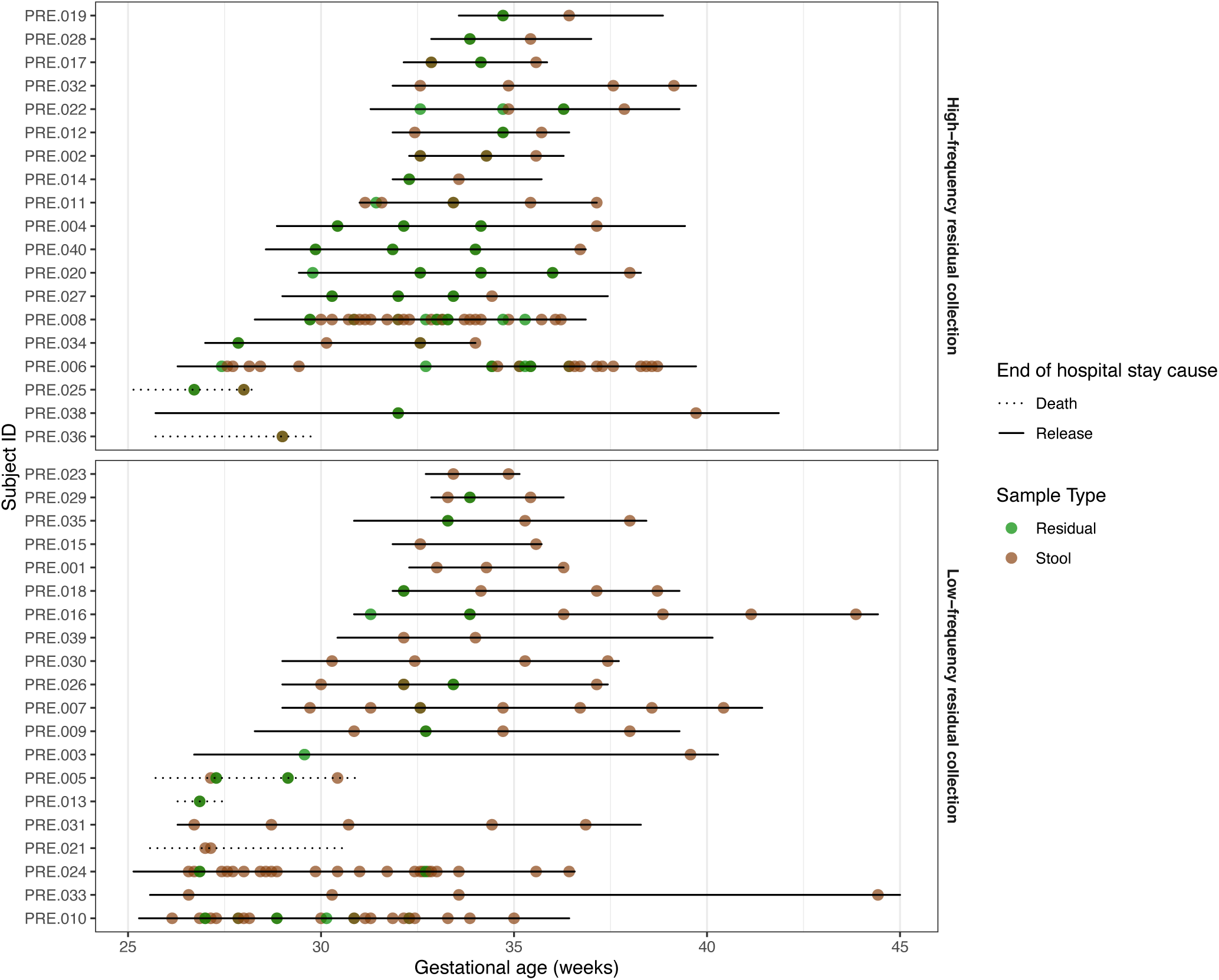
Sample map showing the stool and gastric residuals samples collected along the cohort. Subjects (shown on the y-axis) were divided into 2 groups. Gestational age (in weeks) is shown on the x-axis. The line type represents the clinical outcome of each infant. Dots along the hospitalizations mark samples that were sequenced. Samples (dots) are colored according to their sample type.

**Table 1.** Comparison of clinical metrics across study groups.

| Parameter | Low-frequency residuals group (*n*=20) | High-frequency residuals group (*n*=19) | p-value |
| --- | --- | --- | --- |
| Gestational age at birth | 28w + 5d | 29w + 4d | 0.35 |
| Birthweight (grams) | 1170 | 1130 | 0.76 |
| Length of hospitalization (days) | 60.2 | 50.2 | 0.28 |
| Birth z-score | 0.049 | -0.548 | 0.059 |
| Hospitalization delta z-score | 1.44 | 1.15 | 0.24 |

### Gastric residuals microbiome

To examine the various patterns of the gastric residual (GR) microbiome of preterm infants, we performed metagenomic sequencing on 69 samples from 30 preterm infants. In general, the GR samples often had low-diversity microbial communities, dominated by single-few bacterial species **(Fig. 2A)**, enabling a clean clustering into 11 distinct clusters (**Fig. 2A,B**; using partitioning around means (PAM) algorithm, with Bray-Curtis dissimilarity; **Methods**). Each of the 11 clusters can be characterized by its dominant representative bacteria, commonly from the *Klebsiella*, *Streptococcus* and *Staphylococcus* genera (**Fig. 2A**), known inhabitants of the gut and skin microbial populations^59–67^. The cluster assignment was done irrespective of the GR collection frequency, and we found most clusters to contain samples from both frequencies (**Fig 2C**).

**Figure 2.**
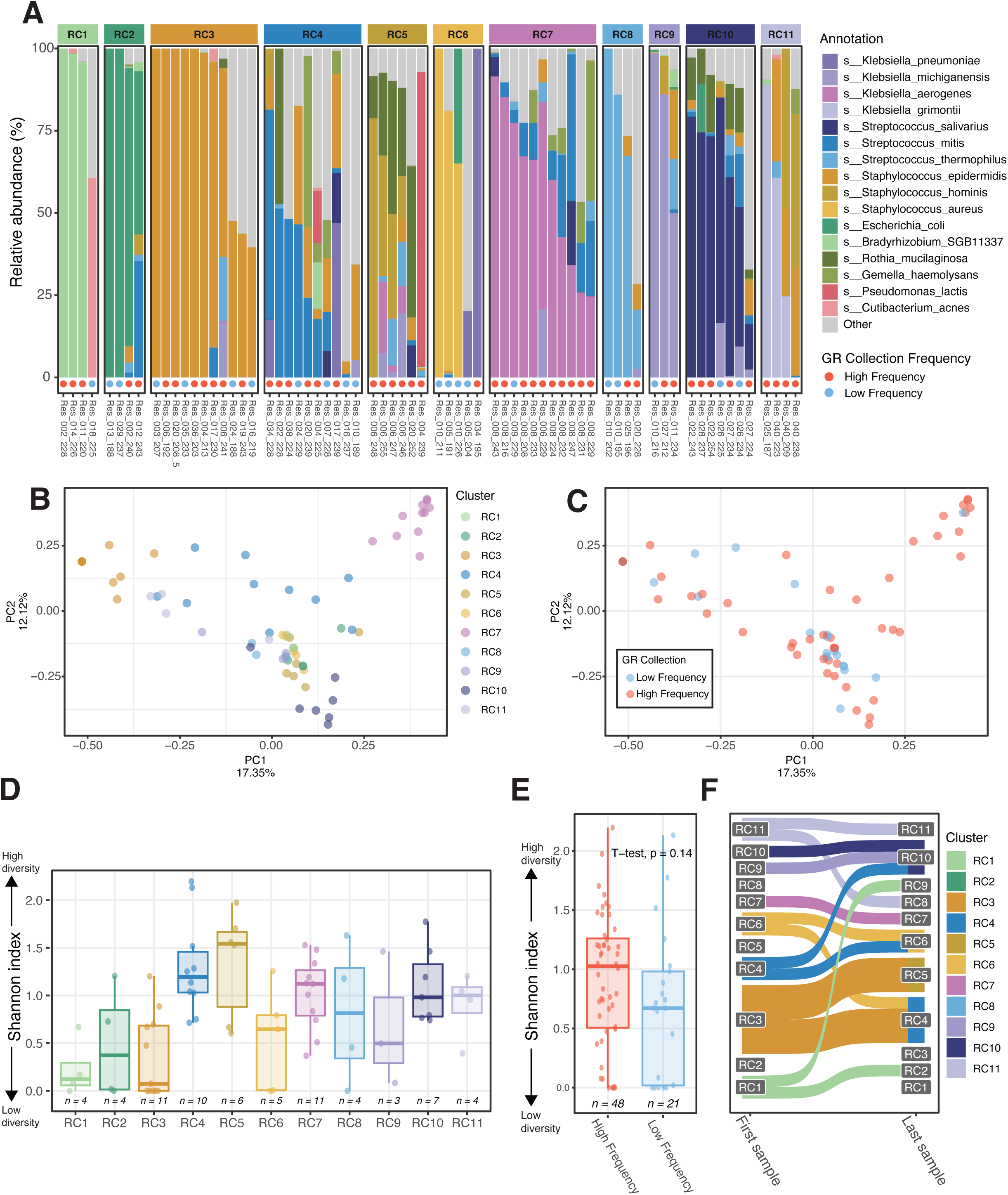
Gastric residuals (GR) microbiome. **A)** Microbial profiles of all GR samples. Each bar represents a single sample. Samples (bars) are separated according to their assigned cluster. Bars are colored according to the taxa. GR collection frequency for the sample subject is represented in a dot under bars. **B,C)** Principal coordinate analysis (PCoA) of GR samples by sample cluster (B) and GR collection frequency (C), respectively. Each sample is represented as a dot on the plot and is colored according to its assigned cluster. The distance between two samples (dots) is proportional to the microbial distance of the two samples. **D, E**) Comparison of the alpha-diversity (community richness) of GR samples between clusters (D) and GR collection frequency (E), respectively. Box represents 25% and 75% quantiles. Each dot represents a sample. Number of samples within each cluster is noted under each box. **F)** Dynamics of early life gastric residuals stability. Lines connecting between the right (first sample) and left (last sample) columns represent a single infant. The width of the lines connecting the columns is proportional to the number of infants sharing this trajectory. Clusters appear as the y-axis and are colored according to their representative bacteria.

Next, we explored the bacterial richness of GR samples, and as the stomach is known for its acidity, we *a priori* expected to find bacteria that thrive in low-pH conditions, such as *Lactobacilli* or *Helicobacter*, which are commonly found in acidic niches such as the vagina, and mature gastric microbiome, respectively^68–72^. In theory, the acidic environment in the stomach could present challenging conditions for bacterial colonization, permitting only a few shared colonization-eligible bacteria, thus resulting in a likely low bacterial richness. To our surprise, *Lactobacillus* species were not abundant in any of the GR samples **(Fig. 2A)**. Most GR samples had low-diversity communities, as reflected by Shannon index **(Fig. 2D)**. When considering individual clusters, we found that clusters RC4, RC5, and RC7, which were dominated by *Streptococcus mitis*, *Staphylococcus hominis*, and *Klebsiella aerogenes*, respectively, had the richest communities (**Fig. 2D**). With respect to GR collection frequency, we found that higher collection frequency tended to have a richer GR community, yet this association was not statistically significant (**Fig. 2E**).

Finally, we assessed the stability of the GR microbiome during the hospitalization period of the infants in the NICU. To understand the intra-subject variance of this developing microbial niche, we selected the first and last GR samples collected per infant and compared the assigned microbial cluster at the beginning and end of hospitalization. We found that clusters RC2, RC5 and RC8, which were rich in *Escherichia coli, Staphylococcus hominis* and *Streptococcus thermophilus*, respectively, were completely missing in the first GR samples (**Fig. 2E**). In contrast, clusters RC1 and RC3, which were rich in *Bradyrhizobium* and *Staphylococcus epidermidis*, respectively, were completely missing in the last GR samples (**Fig. 2E**). Taken together, these dynamics could suggest an age-dependent development in the GR environment where some early colonizers do not sustain in the developing stomach. Additionally, some gastric bacteria may lack the ability to efficiently thrive in a young GR niche while retaining the capacity to colonize it later on in early life.

### Overview of the gut microbiome of preterm infants

First, to get an overview of the premature infant gut microbiome and its development across the hospitalization period (mean 55.3 days, median 55 days), we performed metagenomic sequencing and profiled the microbiome of 132 stool samples from 39 subjects (**Methods**; **Fig 3A**). Overall, the premature infants’ gut microbiome maintained low bacterial diversity, naturally falling into 8 distinct clusters (generated using partitioning around medioids (PAM) analysis with Bray-Curtis dissimilarity; **Methods**, **Fig 3B**). Each cluster was characterized by a dominant representative bacteria, mainly from the *Enterobacteriaceae* family, such as *Escherichia coli* and *Klebsiella spp* **(Fig 3A)**. As expected, some of the cluster-representative taxa were previously reported as popular colonizers, and potentially pathobionts, of the premature infant gut, such as members of the *Staphylococcus, Klebsiella, Enterococcus* and *Escherichia* genera^52,73–78^. We found the clustering classification to be independent of the mode of delivery, age at sampling, and the frequency of gastric residuals sampling (**Supp Fig 1A,B,C**). Interestingly, while probiotics are not administered in this NICU, we found *Bifidobacterium* species in 17 premature infants, which is considered a measure of overall health in early life^18,21,22,24^ (**Supp Fig 2**).

**Figure 3.**
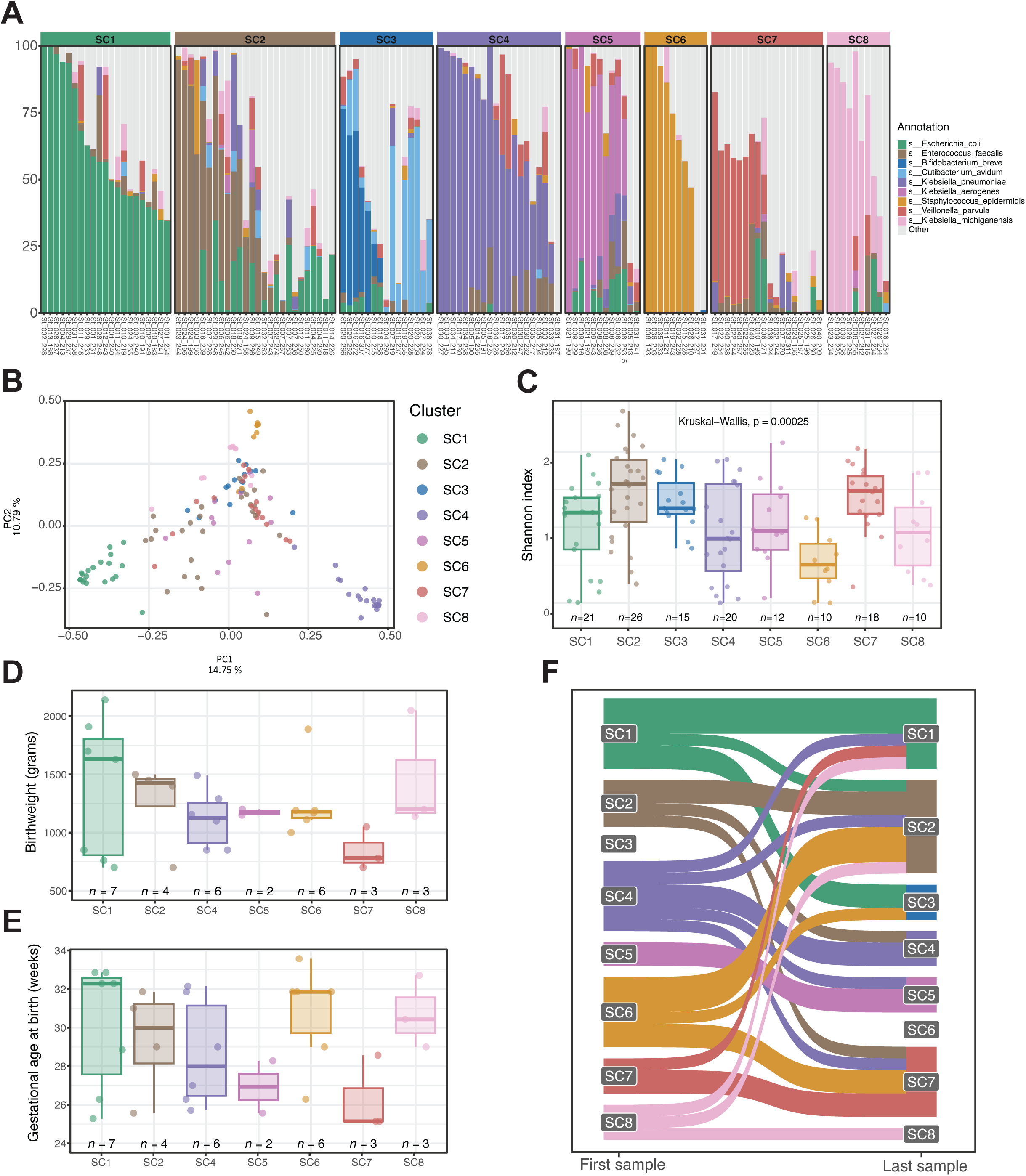
Stool microbiome. **A)** Microbial profiles of 199 stool samples. Each bar represents a single sample. Samples (bars) are separated according to their assigned cluster. Bars are colored according to the taxa. **B)** Principal coordinate analysis (PCoA) of stool samples. Each sample is represented as a dot on the plot and is colored according to its assigned cluster. The distance between two samples (dots) is proportional to the microbial distance of the two samples. **C)** Comparison of the alpha-diversity (community richness) of stool samples between clusters. Box represents 25% and 75% quantiles. Each dot represents a sample. Number of samples within each cluster is noted under each box. **D, E)** Comparison of the birthweight and gestational age at birth, respectively, between clusters of subjects’ first samples. For each infant, the first sample was used to categorize the infant to a specific cluster. Box represents 25% and 75% quantiles. Each dot represents an infant. Number of subjects within each cluster is noted under each box. **F)** Dynamics of early life stool microbiome stability. Lines connecting between the right (first sample) and left (last sample) columns represent a single infant. The width of the lines connecting the columns is proportional to the number of infants sharing this trajectory. Clusters appear as the y-axis and are colored according to the representative bacteria.

Our observation that various bacteria can excel and thrive in the premature infant’s gut raises the question of whether different dominant taxa allow for different microbial community dynamics to develop. To compare the different microbial clusters in terms of community evenness and richness, we calculated the Shannon and the Chao1 indices, respectively. We found that while most of the clusters had similar alpha diversity metrics, SC6, dominated by *Staphylococcus epidermidis*, maintained a markedly decreased alpha diversity in both evenness and species richness **(Fig 3C; Supp Fig 3)**. These findings suggest a colonization of the immature gut environment by *S.epidermidis* resulting in an uneven and low diversity microbial community. Additionally, these findings may hint that *S. epidermidis*, when dominating a niche, challenges other bacteria seeking to thrive alongside it.

### Earliest microbiome profile and its clinical associations

Next, we sought to understand if the assigned microbial cluster can provide insightful information regarding the infant’s health status. To evaluate a possible correlation between the microbial profile and the subject’s anthropometric measurements, we decided to focus on the earliest time point for each infant, collected within the first two weeks of life. We then examined the association of the assigned cluster of the first stool sample with birthweight and gestational age at birth **(Fig 3D, 3E)**. We found that infants in SC6 had an overall advanced gestational age at birth with a relatively low birthweight, (small for gestational age - SGA). In contrast, infants in SC1, dominated by *Escherichia coli,* had the highest median gestational age and weight at birth. Our results highlight a possible correlation between *S. epidermidis* dominated stool samples in early life and birth weight that is inappropriate for their age. To check whether mode of delivery was a confounding factor in our results, we validated that birthweight and gestational age at birth were not associated with mode of delivery **(Supp Fig 4)**.

### Microbiome dynamics during hospitalizations

Premature infants are often hospitalized for a considerable period following birth, allowing us to examine the dynamics of their microbiome during hospitalization. To examine whether the gut microbiome converges towards a consensus community, we compared the microbial composition of the first and last sample of each infant, highlighting shifts in microbial cluster assignments throughout the hospitalization period **(Fig 3F)**. Surprisingly, we found that none of the infants had their last stool sample assigned the *S. epidermidis-*rich SC6 despite it being a relatively prevalent cluster among the first stool samples. Notably we found SC3, a cluster rich in *Bifidobacterium breve*, to be represented across the last stool samples despite being absent in the first samples **(Supp Fig 2)**. Overall, when examining additional time points throughout the hospitalization, we observed that samples annotated as SC6 tended to be collected at earlier time points, whereas samples assigned to SC3 were collected at later stages **(Supp Fig 5)**.

### Shared microbial strains colonize both gastric and gut environment

First, we compared the overall microbial composition in the two environments, to assess the impact of the different niche settings on their residing microbiome. Using a principal coordinate analysis with the Bray-Curtis dissimilarity metric, we found no significant separation between the two sample types, implying the presence of similar bacteria across the two niches **(Fig 4A)**. Specifically, we identified *Staphylococcus epidermidis, Escherichia coli* and *Klebsiella pneumoniae* to maintain a pivotal role in explaining the variability across these samples **(Fig 4B,C,D)**.

**Figure 4.**
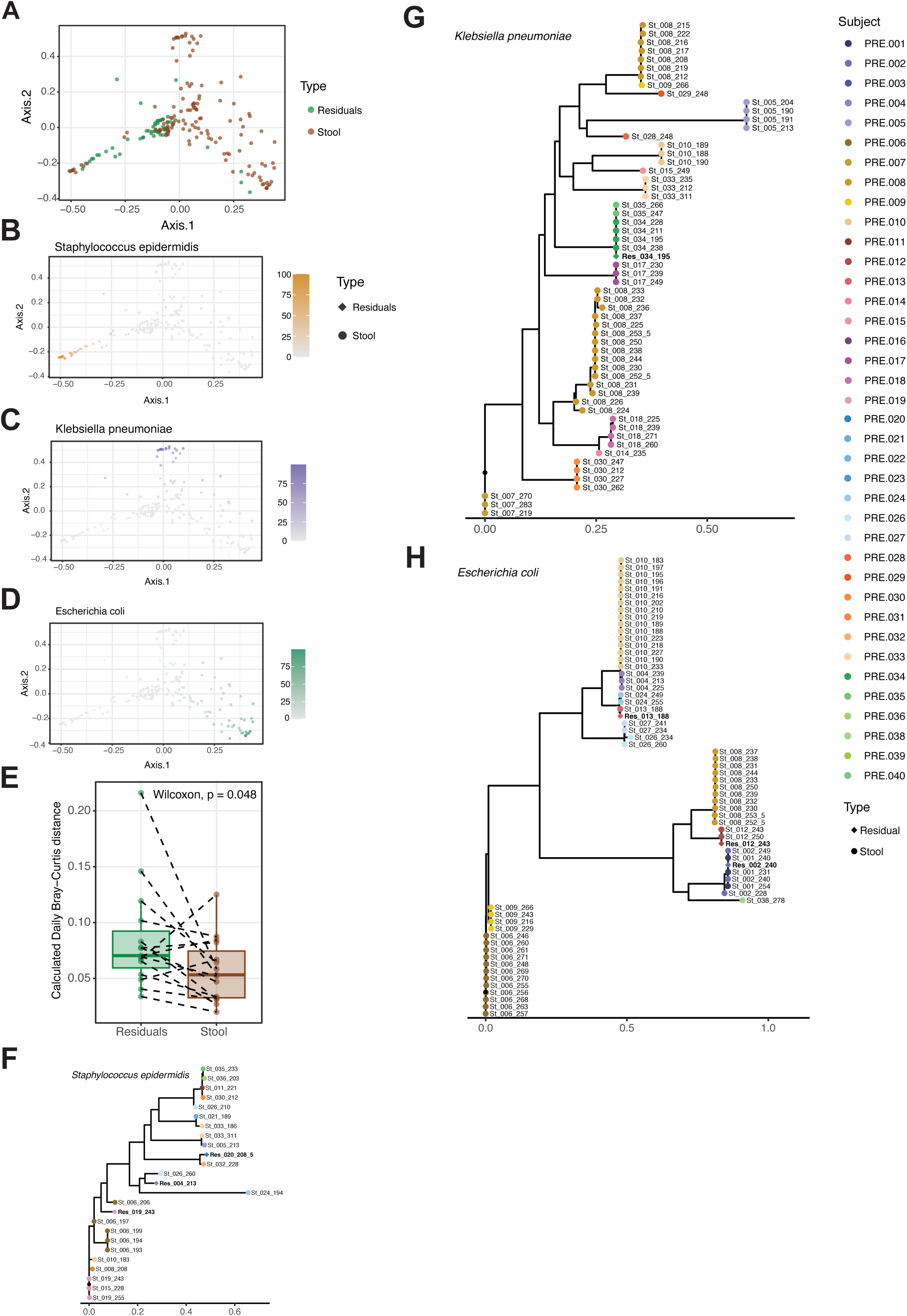
Stool and gastric residuals microbiome. **A,B,C,D)** Principal coordinate analysis (PCoA) of stool and GR samples by sample type (A) and *S.epidermidis* (B), *K.pneumoniae* (C), and *E.coli* (D) abundance. Each sample is represented as a dot on the plot. The distance between two samples (dots) is proportional to the microbial distance of the two samples. **E)** Comparison by sample type of average microbial dissimlarity across two consecutive samples. Each dot represents the average daily Bray-Curtis dissimlarity of a single subject. Measurements (dots) from the same infant are connected with a dashed line. **F,G,H)** Strain level analysis of key taxa. Phylogenetic trees of *S.epidermidis* (F), *K.pneumoniae* (G), and *E.coli* (H). Each leaf (dots) represents a single sample.The phylogenetic distance between dominant strains of two samples is the sum of the distances on the x-axis (branches of the tree). Leafs (samples) are colored by the subject from which the sample was taken. Leafs (samples) are shaped according to sample type.

We next examined whether the GR microbiome is less or more stable than the gut microbiome. We compared the compositional dissimilarity of consecutive GR and consecutive stool samples, using the Bray-Curtis dissimilarity index. Specifically, we looked at consecutive samples that were collected within 3 weeks of each other. Our comparison was conducted across 36 and 78 pairs of GR and stool samples, respectively. We found consecutive GR samples to be less similar to each other compared to stool samples, indicating that the gut microbiome is more stable than the gastric microbial community **(Fig 4E**, *p-value = 0.048, paired Wilcoxon*).

Lastly, we turned to evaluate the strain similarity within and across body niches. We compared the dominant strain of common species across all stool and gastric residual samples, focusing on the eight representative taxa of the stool clusters **(Fig 4F,G,H; Supp Fig 6A,B,C,D)**. We found that most of these key bacteria, including *Escherichia coli, Klebsiella pneumoniae, Enterococcus faecalis* and *Veillonella parvula* maintained a relatively constant dominant strain in infants throughout their hospitalization **(Fig 4F,G,H; Supp Fig 6A,B,C,D)**. In contrast, in some cases, like for *Staphylococcus epidermidis,* similar strains were found across multiple subjects, thus suggesting an environmental mode of acquisition.

### Early metagenomic detection of pathogenic Klebsiella aerogenes strain

Finally, we focused on a single premature infant (PRE.021) who developed *Klebsiella aerogenes* sepsis, ultimately resulting in death. When examining the strains of *K. aerogenes*, another key bacterium in the developing preterm gut microbiome, we found that although similar strains were shared across several subjects, a completely distinct strain was present in PRE.021 **(Fig 5A)**.

**Figure 5.**
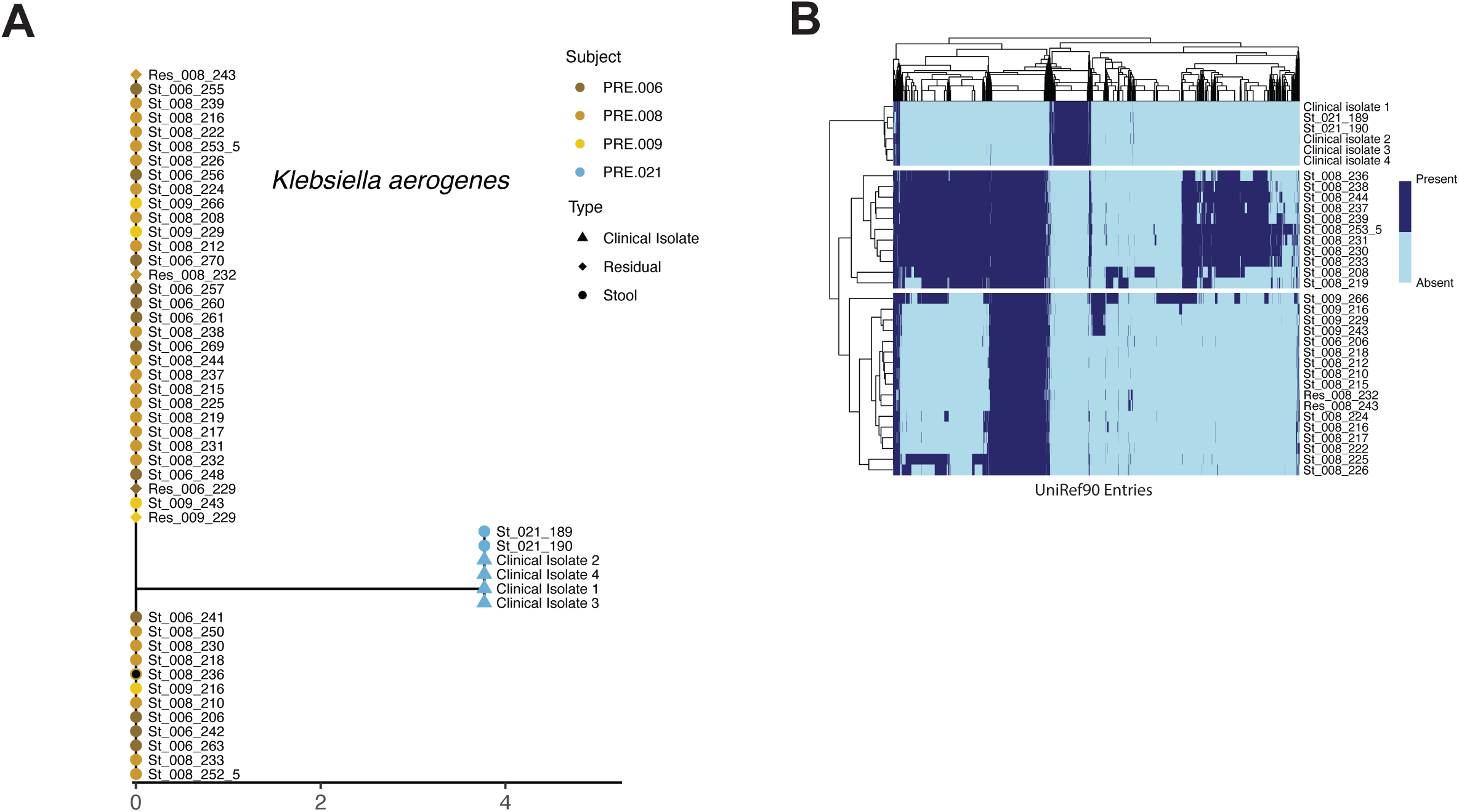
*Klebsiella aerogenes* strain-level comparison. **A)** Phylogenetic tree of *K.aerogenes*. Each leaf (dots) represents a single sample. The phylogenetic distance between dominant strains of two samples is the sum of the distances on the x-axis (branches of the tree). Leafs (samples) are colored by the subject from which the sample was taken. Leafs (samples) are shaped according to sample type. **B)** Heatmap comparing the genomic content of dominant *K.aerogenes* strain across different samples. Each row represents the dominant *K.aerogenes* strain of a single sample or clinical isolate. Each column represents a specific Uniref90 entry. Presence or absence of a Uniref entry within a genome is represented by the fill color. Samples (rows) and Unirefs (columns) are clustered hierarchically. Uniref entries that are shared or absent across all samples are not shown in the plot.

To explore the differences underlying the phylogenetic separation between the dominant *K. aerogenes* strain in PRE.021 and the strains present in other individuals, we carried out a genomic content comparison analysis using PanPhlAn, highlighting 377 and 199 Uniref90 and UniProt entries, respectively, that are found exclusively within PRE.021 and completely missing from strains of other subjects **(Fig 5B)**. We then compared these entries against 12 different *K. aerogenes* reference genomes, using BioCyc^79^, and found 11 genes that were present in the PRE.021 strain and were missing from at least 9 of the 12 reference genomes. These include genes associated with bacterial replication (such as *dnaA, dnaC)*, SoxR regulatory genes (*rsxB)*, catalysing enzymes (*rihC, rfbA, ligB, hrpB)*, iron-sulfur complexes (*iscX*), nitrogen transport (ptsN), and multidrug resistance proteins *(mdtB)*. Specifically, *hpaR* (transcription regulator) and *fabG* (fatty acid elongation) were found to be associated with pathways and gene ontology (GO) groups that are significantly enriched in the *K.aerogenes* strain of PRE.021 compared to reference genomes.

In contrast, we noted 619 and 614, Uniref90 and UniProt entries, respectively, that were consistently missing in the PRE.021 strain, despite being present in all strains of other infants. We found 64 genes to be absent in PRE.021’s *K. aerogenes* yet present in various reference genomes of the bacterium. Of these, *cysS, thrS* (both involved in tRNA functionality), and *fhuA* (iron uptake) were most commonly missing in PRE.021 compared to reference genomes. We did not find a specific pathway or GO to be significantly depleted among *K. aerogenes* of PRE.021 in comparison to references.

Remarkably, we observed this unique strain of *K. aerogenes* in stool seven days prior to its first clinical isolation from PRE.021’s blood culture, despite multiple such culturing efforts throughout this period. This early detection of the pathogenic strain in stool underscores the potential of metagenomic sequencing in the detection of pathogens at an early stage.

## Discussion

In this study, we provide a comprehensive characterization of the gastric residual and stool microbiome of premature infants, revealing distinct microbial community structures and their temporal dynamics during hospitalization. We were able to cluster over 130 stool samples into eight groups based on their bacterial composition, highlighting the varying nature of preterm gut microbiome development across different infants. Despite often being dominated by members of the *Enterobactericeae* family, we also found stool samples to host a high abundance of *Staphylococcus epidermidis*. Samples rich in *S. epidermidis* were mostly prevalent at early time points in the hospitalization and were associated with a lower birthweight for gestational age. Our results reflect the interaction between the gut developmental status and the microbial colonization. We assessed the dynamics of gut microbiome composition across hospitalization and noted that *S. epidermidis-*rich populations were completely missing towards the end of hospitalization. These findings support *S. epidermidis* as a transient colonizer of the premature infant gut and reflect the changes of gut maturation which leads to a less *S. epidermidis*-friendly niche^52^.

In contrast to the dynamic colonization of *S. epidermidis*, we find *Bifidobacterium*-rich communities appearing in late time points of the hospitalization. Higher abundance of *Bifidobacterium* species has been associated with a healthy infant gut^18–21,24,25^, and while these species are commonly found in neonatal probiotic supplements due to their beneficial effect^19,21–23,25^, there was no use of probiotics in our cohort (as instructed by the local ministry of health), thus all colonization reported here was independent of such supplements. We find several *Bifidobacterium* species, notably *B. breve, B. bifidum* and *B. longum* to be present across multiple infants. Our findings reinforce the notion that preterm infant gut development leads towards a more mature and health-associated microbial profile.

To the best of our knowledge, we performed the first metagenomic analysis of the gastric residual (GR) microbiome of preterm infants. Our results suggest variable bacterial colonization patterns and unstable intra-subject microbial profiles in early life. Specifically, we report high abundances of skin and mucosa-associated bacteria such as *Staphylococci* and *Streptococci*^59–61,67^. Additionally, we found a positive correlation between frequent GR aspirations and increased GR microbial diversity. Taken together, our findings suggest a possible translocation of bacteria from the oropharynx to the gastric environment via the nasogastric tube. Furthermore, our findings challenge our initial hypothesis that the stomach environment applies selective pressures on bacterial colonization, permitting only acid-tolerant bacteria to thrive, yet this hypothesis may still hold in term infants with mature gastrointestinal tract. Interestingly, we analyze the microbial trajectory of GR in preterm infants and note shared bacteria throughout the hospitalization period.

One significant finding of our GR microbiome analysis is the notable abundance of a *Bradyrhizobium* species, which is considered to be an environmental bacteria often found in soil^80–82^. We first suspected *Bradyrhizobium* presence to be a result of contamination during the sample preparation, but our contamination analysis did not support this hypothesis **(Methods)**. Within our data, we found *Bradyrhizobium* presence in GR samples of 9 different infants (5 of which also had *Bradyrhizobium* in their stool samples). Combining these results, despite not being able to completely rule out possible contamination originating in the sample collection step or the NICU environment, we conclude that the *Bradyrhizobium* abundance observed genuinely reflects its presence within the premature infant microbiome.

Clinically, our work promotes the potential for early detection of pathogenic bacteria using sequencing approaches. We were able to identify the presence of a pathogenic *Klebsiella aerogenes* in the stool, a full week before its first isolation from clinical specimens. Importantly, clinical sampling was performed in an attempt to monitor a possible infection as early as nine days ahead of our stool sample time point on a near-daily basis, yet it was still undetected.

Our findings offer novel insights into the colonization patterns of the neonatal gastrointestinal tract in two different organs, and importantly present grounds for hope regarding future diagnostic approaches incorporating the sensitivity advantage of metagenomic sequencing.

## Methods

### Study population

Our study consisted of 39 premature infants, born before 37 weeks of gestation and hospitalized in Hadassah Ein Karem Medical Center, Jerusalem. Study participants were randomized into 2 study arms varying by the frequency of gastric residuals (GR) aspiration. Both study groups were exposed to GR collection upon clinical indication, but additionally, in the high-frequency group, sampling also took place before feeding. Samples and clinical data were collected by caregivers at the neonatal intensive care unit. Parents provided informed consent for subject participation. Our study was approved by Hadassah Medical Center’s ethics committee, IRB number 0107-23-HMO.

### Sample collection

Stool and gastric residuals samples were collected as part of the cohort from premature infants from birth and throughout their hospitalization. Study subjects were randomized into 2 groups of varying gastric residuals (GR) collection frequency. Both the low- and high-frequency GR groups underwent GR sampling upon clinical indication, and additionally, in the high-frequency GR group, gastric content was collected before feeding. Stool samples were collected using eSwab® with 1 ml of liquid Amies medium in order to preserve bacterial population. GR were stored in sterile tubes. Both sample types were collected by the neonatal intensive care team at Hadassah Ein Kerem and stored at 4 °C for up to 24 h and then collected to the lab and stored long term at −80°C.

### Metagenomic library construction and sequencing

DNA extraction from stool samples was done using DNeasy PowerSoil Pro Kit (#47014, QIAGEN). We used the Nextera XT DNA Library Preparation kit (FC-131-1096, Illumina) to create Illumina Sequencing libraries, in accordance to the manufacturer’s recommended protocol using half of the volume and the DNA. Using NextSeq 500, we performed single-end 150 bp sequencing of samples.

### Metagenomic analysis

Using Bowtie2 (2.4.5-1)^83^ and an in house pipeline we removed host reads that aligned to the human genome. Samples were filtered and trimmed for Nextera adapters using fastq-mcf, ea-utils ^84^(1.05). Taxonomic profiling was done using MetaPhlAn4^85,86^ with our unique database^87^, on the background of the mpa_vJun23_CHOCOPhlAnSGB_202307 marker set. We carried out strain-level analysis using StrainPhlAn 4^85,86^ with default parameters and -- sample_with_n_markers 70 and --markers_in_n_samples 70. Further analysis was done using an in house R (4.2.2) script utilizing dplyr^88^ (1.1.2), tidyr^89^ (1.3.0) and tidyverse^90^ (2.0.0). Statistical tests comparing between groups were carried out using the *stat_compare_means()* function of the ggpubr^91^ package.Plots were created using ggplot2^92^ (3.4.2), colors were used from RColorBrewer^93^(1.1–3) and pals^94^ (1.7). Heatmaps were created using pheatmap^95^. Alpha and beta diversity were calculated using “diversity” (Shannon index) and “vegdist” (Bray-Curtis dissimilarity) from the vegan^96^ (2.6–4) package and the PCoA was created using the ape^97^ (5.7- 1) package. Phylogenetic tree was produced using ggtree^98^ (3.6.2) and the sankey plots were created using ggsankey^99^ (0.0.99999).

### Statistical analysis

No statistical method was used to predetermine the sample size. The investigators were not blinded to allocation during experiments and outcome assessment. Independent t-test was performed to test between groups when mentioned using the R function “t-test”. Intra-subject paired t-test was done to compare Bray-Curtis dissimilarities of consecutive stool and gastric residuals samples.

### Clustering analysis

Separation of samples into clusters was done using the partitioning around medians (PAM) model. First, the Bray-Curtis dissimilarity was calculated for each pair of samples, representing the compositional distance between the microbial profiles. Next, to figure out the number of clusters for each sample type, we used the *fpc* (2.2-13)package^100^. Lastly, in order to separate the data into clusters, we applied the *pam* function within the *cluster* (2.1.6) package^101^.

### Bradyrhizobium contamination analysis

To check for possible contamination during DNA extraction, library preparation and sequencing resulting in presence of *Bradyrhizobium* we used the R package *decontam*(1.16.0)^102^. Analysis was performed when taking into account the sample extraction and library preparation batches. To validate the results, we also checked for possible contaminations by sample type, thus analysing the stool and gastric residuals separately.

### Gene enrichment and depletion analysis for pathogenic Klebsiella aerogenes

PanPhlAn^103^ was used to compare the genomic content of *Klebsiella aerogenes* across strains found in our cohort as well as reference genomes. For the enrichment, we selected Uniref90 entries that were consistently present in all data of PRE.021 but completely absent from strains of other cohort subjects. Similarly, for the depletion analysis, we selected all the Uniref90 entries that were missing throughout the data of PRE.021 but were always present in strains of other study participants. Next, we used UniProt^104^ ID Mapping online tool to convert Uniref entries into genes. Using BioCyc^79^ we compared these gene lists to 12 reference genomes of *K.aerogenes,* including EA1509E, NCTC418, FDAARGOS_363, FDAARGOS_513, KCTC2190, NCTC9735, NCTC9997, NCTC9793, NCTC8846, NCTC9652, NCTC9667, and NCTC9668. Finally, enrichment and depletion analysis were carried out searching for enriched and depleted genes for pathways, transcriptional/translational regulators, and GO.

### Stool samples subsampling

Within our sequenced stool samples, subjects PRE.006, PRE.008, PRE.010, and PRE.024 had a greater representation. In order to account for this and avoid biased results stemming we subsampled the number of samples from these infants to be included in the analysis. Briefly, we calculated the floor value of the 95th percentile of the number of samples across other study participants. We then selected this number of samples from the over-represented infants, maintaining the first and last sample collected during the hospitalization.

## Acknowledgements and funding

We thank the participants and their families for joining the research and Dr Abed Nasereddin & Dr Idit Shiff from the Genomics Applications Laboratory at the Faculty of Medicine of the Hebrew University for their support with DNA sequencing. Prof Jacob Strahilevitz and Prof Jacob Moran Gilad from the Clinical Microbiology Department at Hadassah Medical Center for providing the clinical strain isolates. This study was funded in part by the Israel Precision Medicine Partnership (IPMP), grant number 3061/22 and by the Azrieli Family Foundation (Faculty fellowship for MY). MY holds the Rosalind, Paul and Robin Berlin Faculty Development Chair in Perinatal Research.

## Author contributions

M.Y. and N.O.S. conceptualized the study. N.O.S and S.E.F. recruited study participants and gained informed consent from the parents of study subjects. N.M., E.H., N.R., and S.S. transferred samples from the hospital to the lab storage. N.M., N.R. and S.S. were in charge of sample DNA extraction and library sequencing. N.M performed all computational analysis. L.J. and S.S. cultured and isolated strains used for this work. M.Y., N.M., and N.O.S. wrote the manuscript with contributions from all other authors.

## Notes

### Competing Interest Statement

The authors have declared no competing interest.

