## Supplemental Figures for "Development of the preterm infant gut and gastric residuals microbiome"

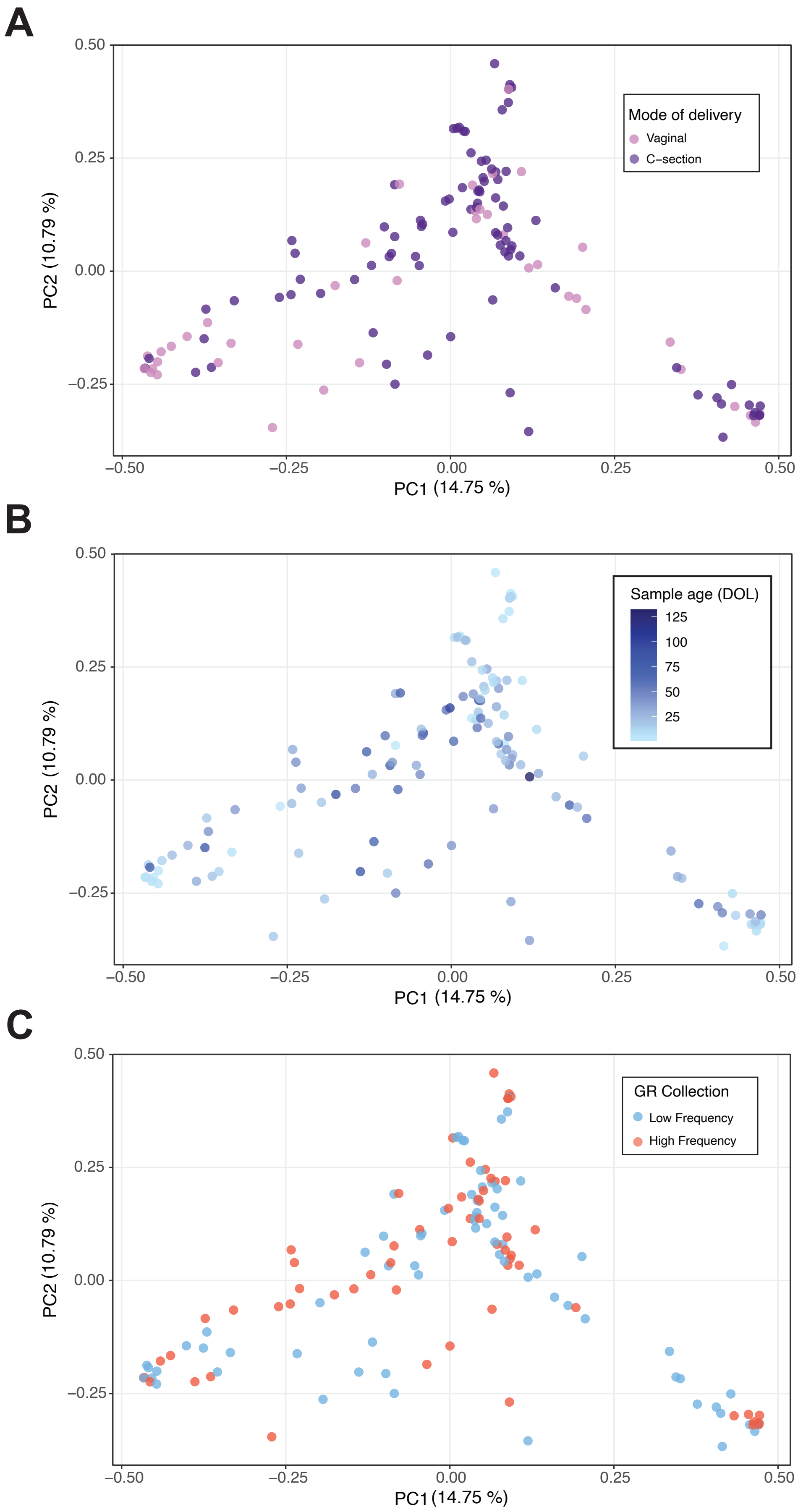


**Supplementary Figure 1**
Principal coordinate analysis (PCoA) of stool samples. Each sample is represented as a dot on the plot and is colored according to the infant's mode of delivery **(A)**, age at time of sample collection **(B)**, and gastric residuals collection frequency **(C)**. The distance between two samples (dots) is proportional to the microbial distance of the two samples.

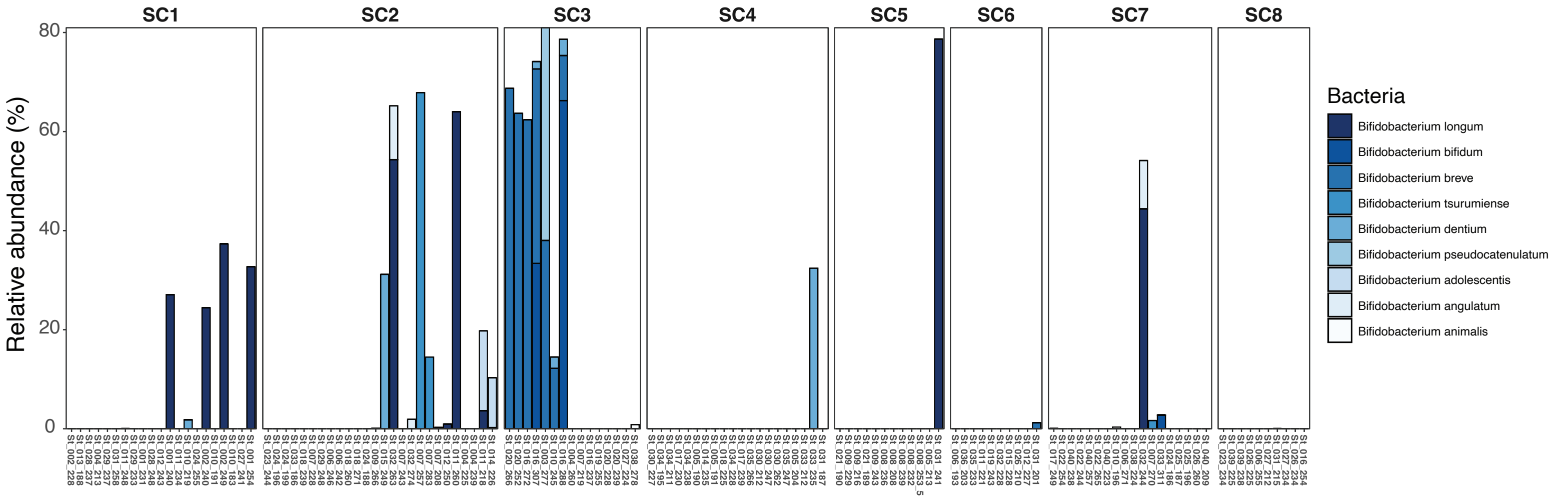


**Supplementary Figure 2**

Microbial profiles of 199 stool samples highlighting Bifidobacterium abundance throughout stool data. Each bar represents a single sample. Samples (bars) are separated according to their assigned cluster. Bars are colored according to the Bifidobacterium species present.

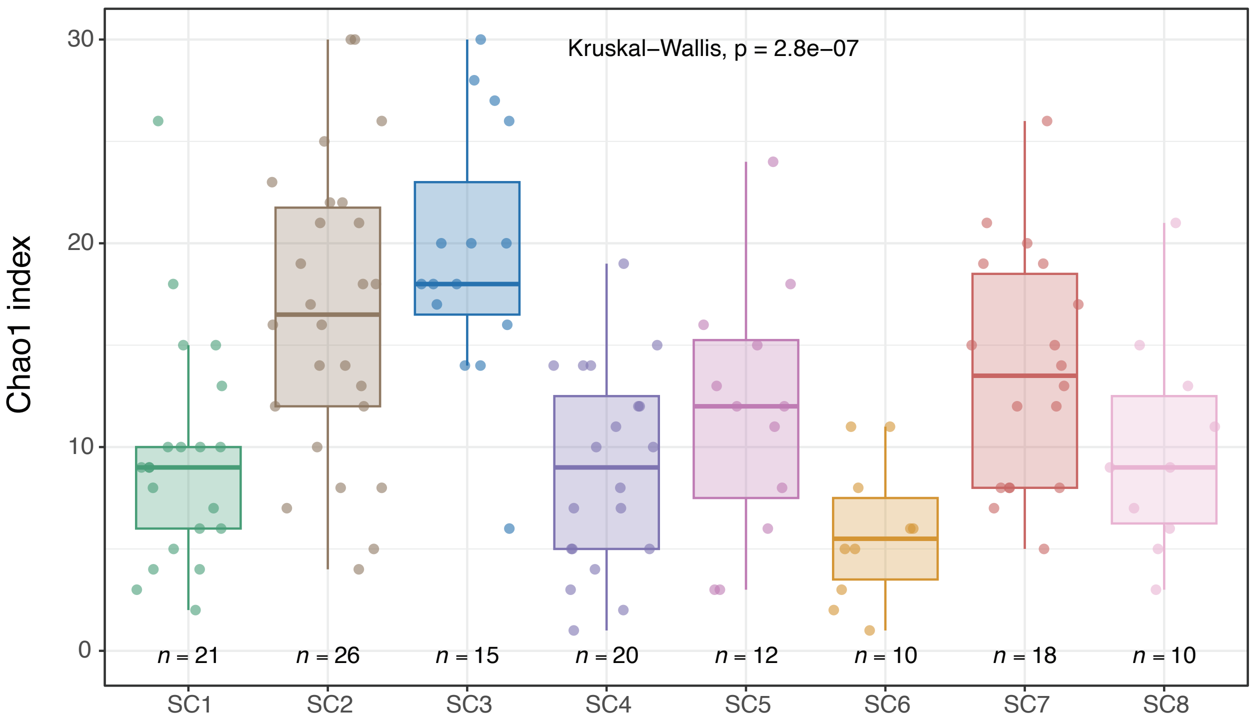


**Supplementary Figure 3**
Comparison of the Chao1 index (community richness) of stool samples between clusters. Box represents 25% and 75% quantiles. Each dot represents a sample, and the number of samples within each cluster is noted under each box.

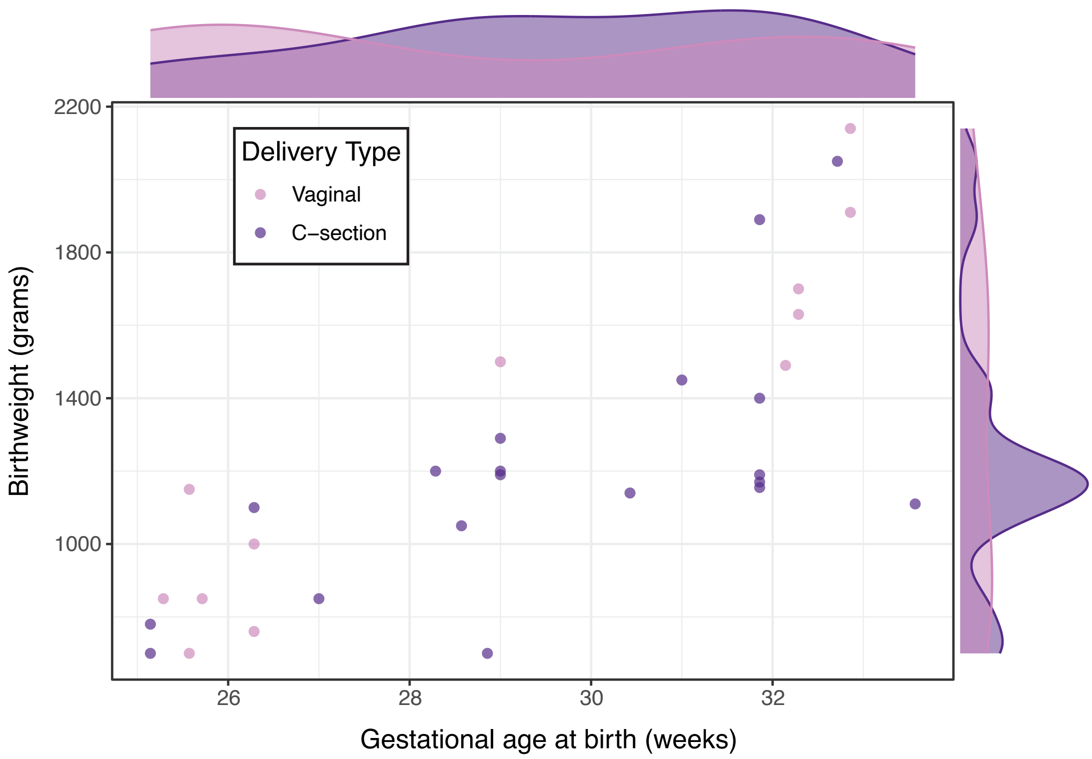


**Supplementary Figure 4**
Anthropometrics at birth across different modes of delivery. Scatterplot showing the gestational age (x-axis) and weight (y-axis) at birth. Each dot represents an individual subject. Infants (dots) are colored according to the mode of delivery. Density plots highlighting the distribution of the data are presented under the y-axis and to the left of the x-axis.

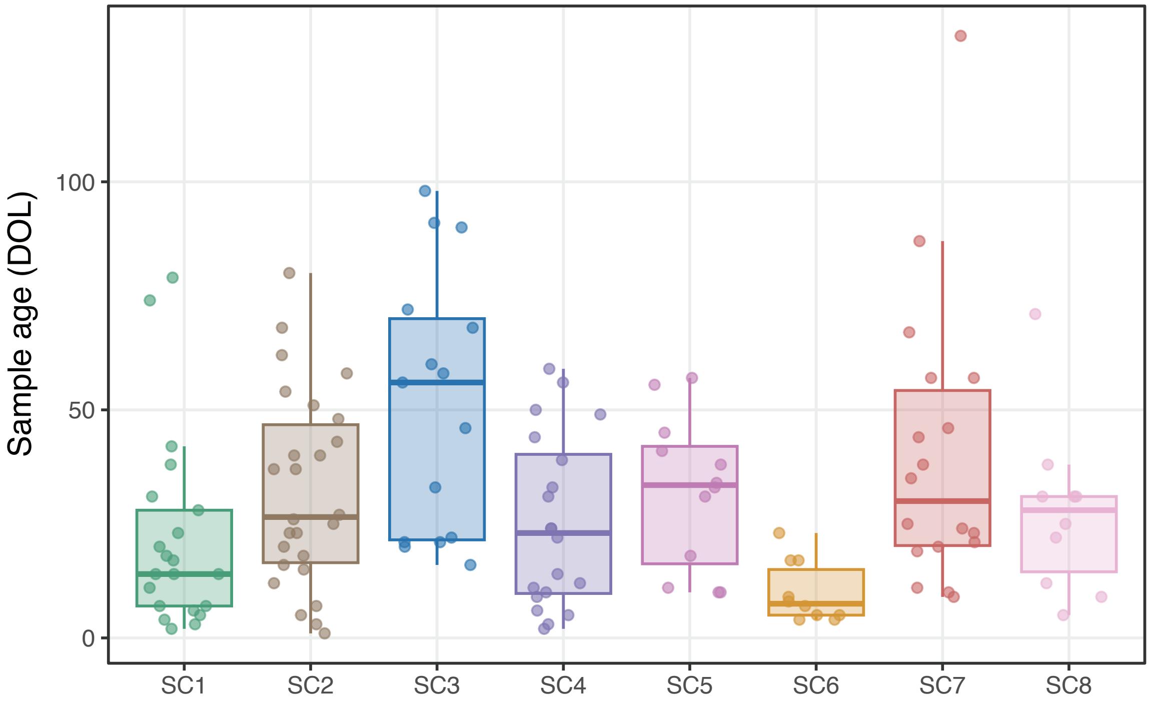


**Supplementary Figure 5**
Comparison of infant age on the day of stool sample collection between clusters.
Box represents 25% and 75% quantiles. Each dot represents a sample.
Dots and boxes are colored according to the assigned microbial cluster.

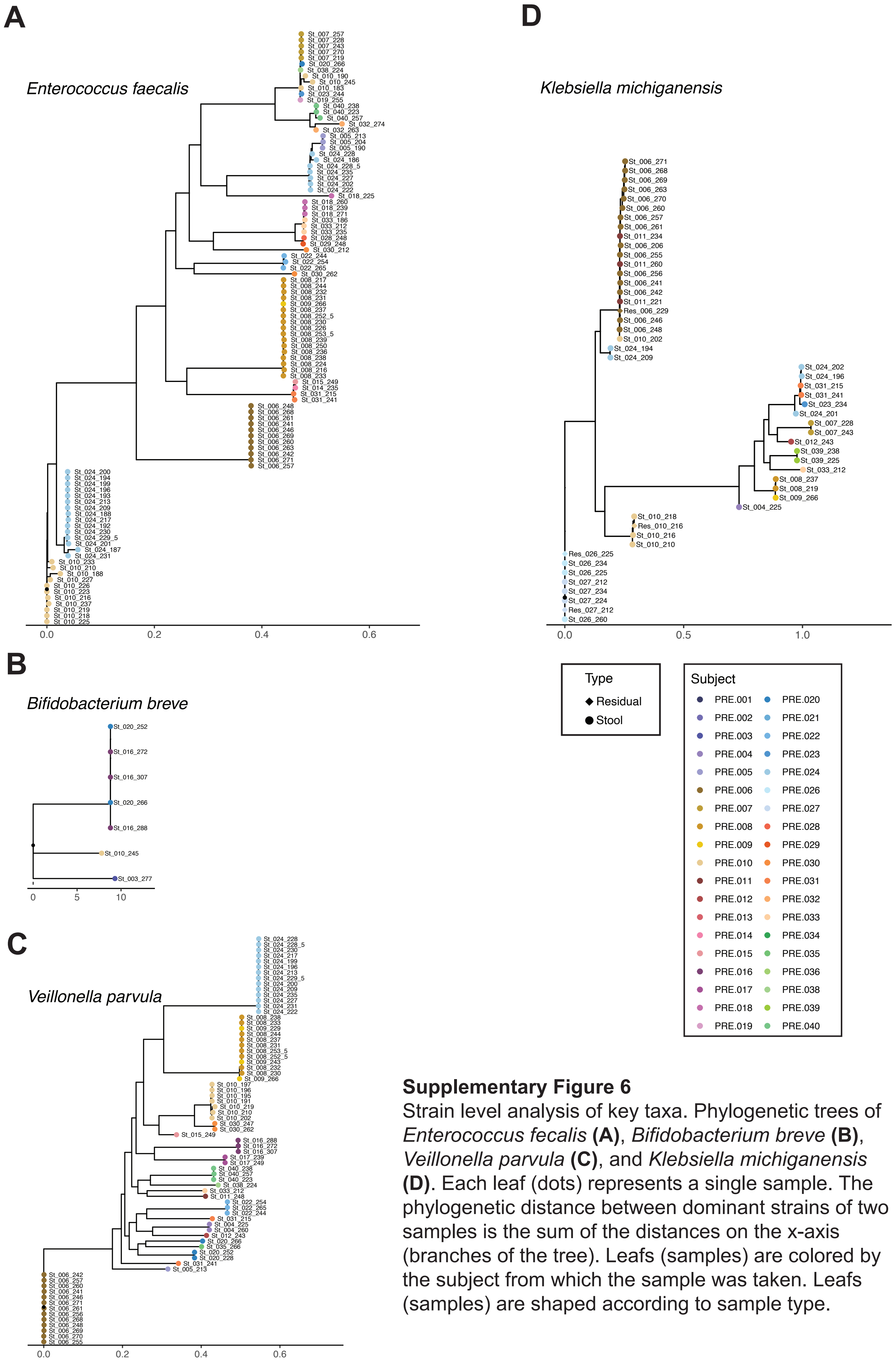


**Supplementary Figure 6**
Strain level analysis of key taxa. Phylogenetic trees of *Enterococcus fecalis* **(A)**, *Bifidobacterium breve* **(B)**, *Veillonella parvula* **(C)**, and *Klebsiella michiganensis* **(D)**. Each leaf (dots) represents a single sample. The phylogenetic distance between dominant strains of two samples is the sum of the distances on the x-axis (branches of the tree). Leafs (samples) are colored by the subject from which the sample was taken. Leafs (samples) are shaped according to sample type.
